# Identification of tubulin polymerization inhibitors with a CRISPR-edited cell line with endogenous fluorescent tagging of β-tubulin and Histone 1

**DOI:** 10.1101/2022.09.14.507662

**Authors:** Harutyun Khachatryan, Carlos A. Barrero, John Gordon, Bartlomiej Olszowy, Oscar Perez-Leal

**Affiliations:** Department of Pharmaceutical Sciences, Moulder Center for Drug Discovery, School of Pharmacy, Temple University, Philadelphia, PA

**Keywords:** CRISPR, tubulin, polymerization, inhibitors, high-content, imaging, screening, high-throughput, homologous, recombination, gene, tagging

## Abstract

Tubulin is an essential protein to maintain the cellular structure and for the cell division process. Inhibiting tubulin polymerization has proven to be an effective method for slowing cancer cell growth. Traditionally, identifying tubulin polymerization inhibitors involved using pure tubulin for in vitro assays or procedures using cells that require cell fixing and anti-tubulin antibody staining. This study explores using a cell line developed via CRISPR genome editing as a cell model to identify tubulin polymerization inhibitors with live cells without using exogenous staining. The cell line has endogenous tagging with fluorescent proteins of β-tubulin and a nuclear protein to facilitate image cellular segmentation by high-content imaging analysis (HCI). The cells were treated with known tubulin polymerization inhibitors, colchicine and vincristine, and the presence of phenotypic changes that indicate tubulin polymerization inhibition were confirmed via HCI. A library of 429 kinase inhibitors was screened to discover tubulin polymerization inhibitors and three compounds that inhibit tubulin polymerization were found (ON-01910, HMN-214, and KX2-391). Live cell tracking analysis confirms that depolymerization of tubulin occurs rapidly after compound treatments. These results suggest that CRISPR-edited cells with fluorescent endogenous tagging of β-tubulin can be used to screen larger compound libraries containing diverse chemical families to identify novel tubulin polymerization inhibitors.

## INTRODUCTION

Cancer constitutes the second most common cause of death following cardiovascular diseases worldwide [1]. Despite significant progress in the understanding, diagnosis, prevention, and treatment of cancer, it remains a major threat to global human health. As the world’s population continues to expand and age, the World Health Organization estimates that 13.2 million cancer-related deaths will occur worldwide by 2030, compared to 7.6 million in 2008 [2].

Of the many approaches used to treat cancer, chemotherapy is one of the most common and effective tools used by oncologists [3]. In chemotherapy, drugs that disrupt microtubule/tubulin dynamics are commonly employed [4]. These drugs directly interfere with the cell’s microtubule system as opposed to acting on the DNA [5]. Microtubules are cytoskeletal structures that allow a cell to maintain its shape. They are also critical in mitotic chromosome separation and create the mitotic spindle, and are thus essential for the successful completion of cell division [6].

Microtubules are composed of primarily alpha and β-tubulin subunits assembled into linear protofilaments that wind together to form a hollow cylinder [7]. These structures can rapidly grow via polymerization and shrink via depolymerization [8]. Epipodophyllotoxins, taxanes, vinca alkaloids, epitholones, and macrolides are among the most common chemotherapeutic drugs that impair microtubule activity [9]. Microtubule targeting agents work by interfering with the dynamics of the microtubule, either inhibiting polymerization or depolymerization.

Microtubule targeting agents interact with tubulin via at least four different binding sites: laulimalide, taxane/epothilone, vinca alkaloid, and colchicine [10]. Polymerization inhibitors mainly act on the vinca alkaloid and colchicine binding sites, while depolymerization inhibitors act on the taxane binding site [11]. Vinca alkaloids, unlike colchicine, attach directly to the microtubule. They do not form a complex with soluble tubulin or copolymerize to form the microtubule, but they are capable of causing a conformational shift in tubulin in the context of tubulin self-association [12].

While tubulin polymerization inhibitors have made great progress in the treatment of many forms of cancer, they have several drawbacks, including hematopoietic and neurologic toxicities, inconvenience formulation, and resistance. Therefore, identifying novel tubulin polymerization inhibitors is necessary to combat these barriers [9].

Traditionally, compounds that are inhibitors of tubulin polymerization have been found using pure tubulin in vitro [13] or procedures that require immunostaining of cells [14], thus limiting the ability to study dynamic subcellular changes. We recently reported a new technology that facilitates using CRISPR genome editing for developing cell lines called FAST-HDR [15]. This technology allows the generation of cell lines by tagging multiple endogenous genes with fluorescent proteins. This approach facilitates studying endogenous proteins in live cells without using cell fixing, immunostaining or recombinant protein overexpression [15].

This work demonstrates the use of a cell line with endogenous tagging with fluorescent proteins of β-tubulin and histone 1 to identify novel tubulin polymerization inhibitors. We screened a library of kinase inhibitors and identified and characterized three compounds as tubulin polymerization inhibitors by using high-content imaging (HCI) analysis of live cells.

## MATERIALS AND METHODS

### Cell line development

HeLa cells with endogenous tagging with fluorescent proteins of β-tubulin (mClover3), histone 1 (mTagBFP2), and p62-SQSTM1 (mRuby3) were developed with CRISPR and the FAST-HDR vector system, and were recently described [15]. These cells are commercially available from ExpressCells, Inc. A clonal cell line was derived by single cell sorting with the Hana Single Cell Sorter Instrument (Namocell) by using the green fluorescent channel and sorting into a 96 well plate. Single cell clones were analyzed with a Spark Cyto plate reader (Tecan) by using whole-well imaging for selecting clones derived from a single cell. A clone with uniform and bright β-tubulin fluorescence was selected to expand a cell line for this study.

### High-content imaging and compound screening

The CRISPR-edited HeLa cell line described above was used to screen a small compound library of 429 kinase inhibitors (SelleckChem, Houston, Texas, USA) for high-content imaging drug screening. 384-well CellCarrier Ultra plates (PerkinElmer, Waltham, Massachusetts, USA) were seeded with 5 × 10^3^ cells per well and 16 hours later were treated with the library compounds at a final concentration of 1 uM by using the PerkinElmer JANUS Automated Workstation. The compounds’ effects were assessed 24 hours later with an Operetta CLS Confocal High Content Imaging System (PerkinElmer, Waltham, Massachusetts, USA) and the following parameters: excitation and emission filter combinations: mTagBFP2 (Ex-405, Em-440), mClover3 (Ex-480, Em-513); acquisition time: 500 ms per channel. Two images (with two different channels) were acquired with the same coordinates for each well by using a 40X water immersion objective. As negative controls 32 wells in each 384-well plate were only treated with dimethyl sulfoxide (DMSO) at 1% final concentration. The positive control compounds (colchicine and vincristine) as well as the identified hits in this study (ON-01910, KX2-391, and HMN-214) were acquired for further validation from Cayman Chemicals in powder form and dissolved with DMSO.

### Live-cell confocal imaging analysis

The Operetta CLS Confocal High Content Imaging System (PerkinElmer, Waltham, Massachusetts, USA) was used for live cell image acquisition. The fluorescent proteins were detected with the following excitation and emission filter combinations: mTagBFP2 (Ex-405, Em-440), mClover3 (Ex-480, Em-513). Images were obtained with a 40× water immersion objective and 500 ms of acquisition time per fluorescent channel. All confocal images in this work were acquired with live cells grown in Dulbecco’s Modified Eagle Medium (DMEM) without phenol red (Thermo Fisher Scientific, Waltham, Massachusetts, USA) supplemented with 10% Fetal Bovine Serum (100 uL) in 384-well CellCarrier Ultra plates (PerkinElmer, Waltham, Massachusetts, USA). For time-lapse microscopy, cells were kept under environmentally controlled conditions (37 °C, 5% CO_2_), and images were acquired every three minutes. All conditions were analyzed in triplicates, and the experiments were repeated three independent times. All compounds under study were administered 30 minutes after the start of image acquisition in a concentration range between 1 nM and 4 uM.

### Molecular docking

We used the x-ray diffraction crystallography protein structure of β-tubulin from *bos taurus* (protein data bank code: 4O2B) for molecular docking experiments. This *bos taurus* tubulin isoform has a 100% homology to human β-tubulin (TUBB). The molecular docking was performed with CB-Dock version 2, a cavity detection-guided protein-ligand blind docking web server (Liu Y, 2022) that uses Autodock Vina (version 1.1.2). The SDF structure files of the tested compounds (colchicine, vinblastine, vincristine, paclitaxel, ON-01910, KX2-391, and HMN-214) were downloaded from PubChem. The molecular docking was performed by uploading the 3D structure PDB file of β-tubulin into the server with the SDF file of each compound. For analysis, we selected the docking poses with the strongest Vina score. The generated PDB files of the molecular docking of each compound were visualized with the Maestro software (Schrödinger, Inc., New York, New York, USA).

### Reproducibility and statistical analysis

All figures in this work are representative of at least three independent experiments. The comparison of multiple groups was performed by using one-way analysis of variance with Dunnett’s test within GraphPad Prism (version 9) (GraphPad Software, La Jolla, California, USA); p values < 0.05 were considered statistically significant. Differences between control and treated samples in time-dependent experiments were performed by using two-way analysis of variance within GraphPad Prism.

## RESULTS

### Detecting the inhibition of tubulin polymerization in live cells without using antibodies or chemical staining

The analysis of phenotypic changes caused by compounds that alter tubulin polymerization usually requires the use of antibodies, chemical staining, or promoting the overexpression of fluorescent tubulin [16]. However, some of these techniques prevent the analysis of live cells to evaluate changes that happen over time or can induce microscopical artifacts by uncontrolled overexpression. As an alternative approach, here we explored using a recently developed HeLa cell line with endogenous tagging of native proteins with fluorescent proteins via CRISPR-Cas9 [15]. In this cell line, endogenous histone 1 is tagged with a blue fluorescent protein (mTagBFP2) and allows the detection of the nucleus. Additionally, β-tubulin (TUBB) is tagged with a green fluorescent protein (mClover3) and facilitates the identification of the microtubules in the cytoplasm. In order to evaluate the potential of this cell line for detecting phenotypic changes in tubulin polymerization by HCI analysis, an analysis routine with the Harmony software (version 4.8) was developed. The routine was set up to detect individual cells in each acquired field by analyzing two independent fluorescent channels. In the blue channel, the software detects the nuclei, where a nuclear protein is labeled with a blue fluorescent protein, by using the Find Nuclei algorithm. In the green channel, the software defines the cytoplasm boundaries of every nuclei with the Find Cytoplasm algorithm by detecting tubulin that is labeled with a green fluorescent protein. Once the software identifies each cell, it can analyze texture properties of the cytoplasm of every cell (Figure 1A).

**Figure 1.**
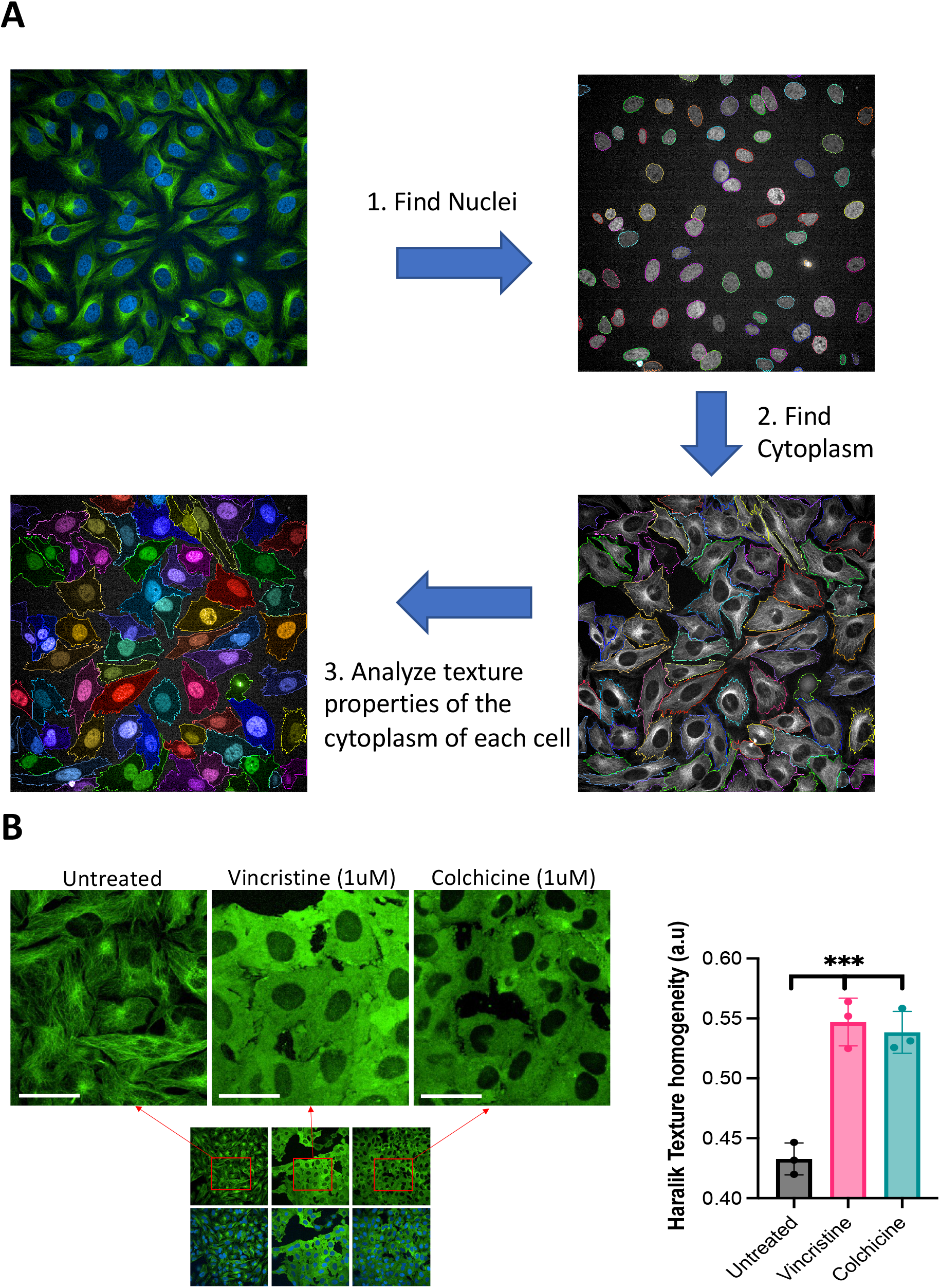
Detection of tubulin polymerization changes in live cells with HCI analysis. (A) The changes in tubulin polymerization were detected and quantified in each microscopic image with the following image analysis workflow: first, detection of the nuclei of every cell in the image by using the blue fluorescent channel for detecting histone 1-mTagBFP2. Second, detection of the cytoplasm of every nuclei that was identified in the first step by using the green fluorescent channel for detecting β-tubulin-mClover3. Third, quantitate the image texture properties of the cytoplasm of every cell. (B) Confocal microscopy images of live CRISPR-edited HeLa cells with fluorescent tagging of histone 1 and β-tubulin. The tubulin polymerization inhibition was detected after treatment with known tubulin polymerization inhibitors (vincristine and colchicine). The changes in tubulin polymerization with each treatment were quantified with the Haralick texture homogeneity algorithm and are shown on the right panel. Differences were analyzed by one-way analysis of variance (ANOVA), and *p* < 0.05 was considered statistically significant. *** *p* < 0.001; error bars represent ± standard deviation (*SD*) of images from triplicate wells. This data is representative of three independent experiments; scale bar = 15 um.

**Figure 2.**
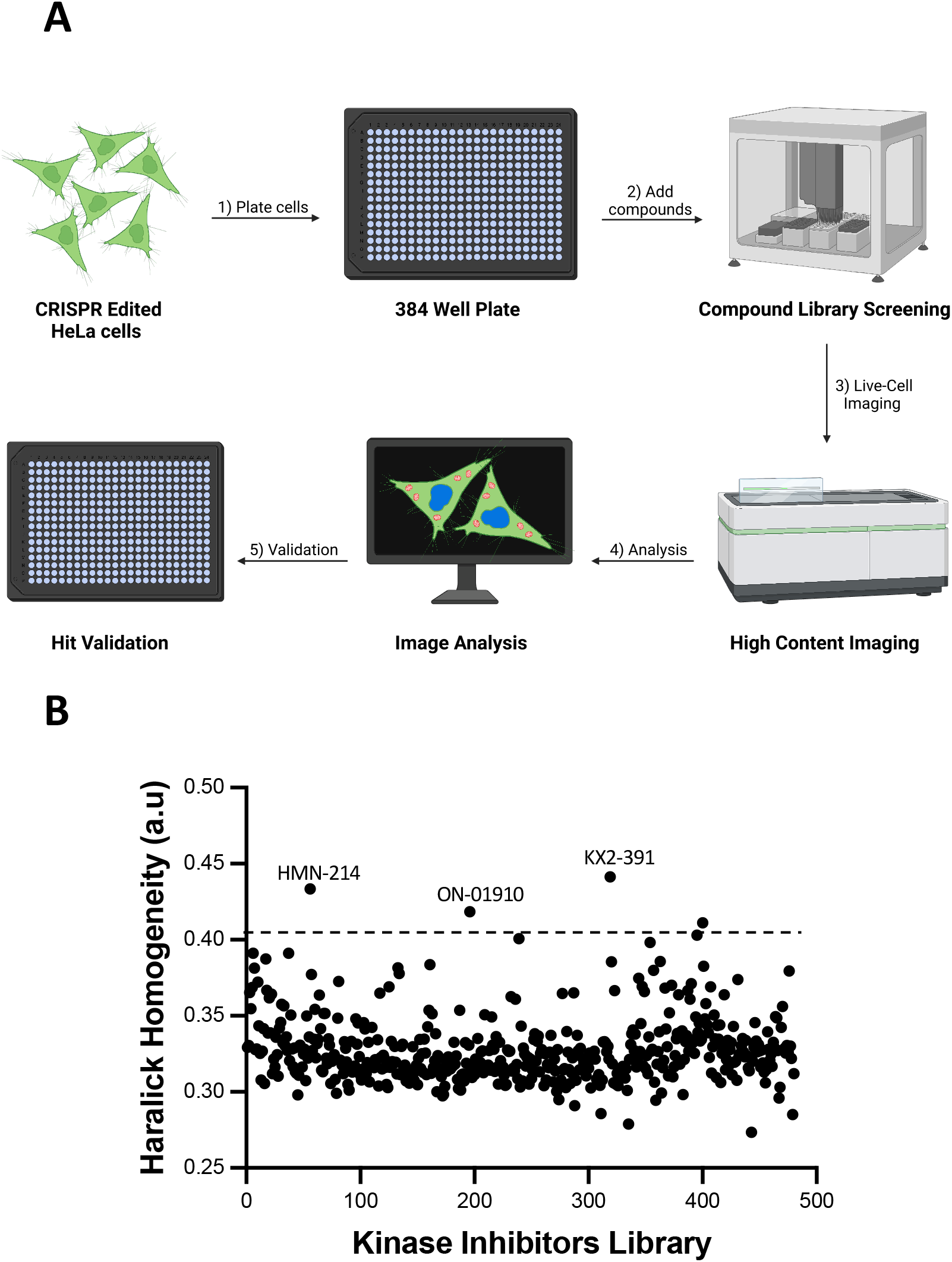
Kinase inhibitor library screening with HCI analysis of live cells. (A) Diagram of the steps required to detect phenotypic changes of CRISPR-edited HeLa cells after treatment with a library of kinase inhibitors. The cells are plated into 384 glass-bottom wells and treated with the compounds of the library for 16 hours. Following a 24-hour incubation period, the cells are imaged directly without cell fixing or immunostaining. The images are analyzed to detect changes in the Haralick homogeneity values as described in Figure 1A. Lastly, the positive hits are further validated. (B) Changes in the Haralick texture homogeneity values of cells treated with a library of 429 kinase inhibitors. Compounds that change the Haralick texture homogeneity higher than the mean of all the compounds plus three standard deviations were considered positive hits for further evaluation. Three compounds (ON-01910, KX2-391, and HMN-214) were manually checked and selected for validation. This data is representative of three independent experiments.

It was then identified how these cells respond to tubulin polymerization inhibitors by first treating the cells with two different tubulin polymerization inhibitors: vincristine and colchicine. Phenotypic changes were identified, where the normal pattern of the microtubules disappears in treated cells (Figure 1B). Next, it was evaluated if these phenotypic changes translate to changes in the texture of the image of tubulin for each cell for automatic detection during compound screening. For that the Haralick texture homogeneity algorithm was used, which is part of the Harmony software analysis package. This algorithm analyzes the homogeneity of the pixels in an image to determine regularity [17]. The analysis of the images from cells treated with known tubulin polymerization inhibitors or controls shows that the Haralick texture homogeneity values increase for vincristine and colchicine, and these changes were statistically significant (Figure 1B).

### Identifying tubulin polymerization inhibitors in a library of kinase inhibitors

The HeLa cell line with endogenous tagging of histone 1 and β-tubulin was used for screening a library of 429 kinase inhibitors to identify tubulin polymerization inhibitors by using HCI analysis. The cells were plated onto 384 well plates, treated with the library of kinase inhibitors for 16 hours, and directly analyzed with an automated high-content imager. The obtained images were analyzed with the procedure depicted in Figure 1A. The Haralick texture homogeneity values of cells treated with all the compounds were determined to identify potential inhibitors of tubulin polymerization. One point of interest were compounds that could induce changes in the Haralick texture homogeneity values that were higher than the mean plus three times the standard deviation of all the compounds. According to the data, four compounds were identified as positive hits; however, after manual inspection of the images of these four wells, only three compounds (ON-01910, HMN-214, and KX2-391) were found to be positive hits.

### Validating kinase inhibitors as tubulin polymerization inhibitors by using live-cell imaging analysis

To confirm that ON-01910, HMN-214, and KX2-391 are inhibitors of tubulin polymerization, a dose response analysis was performed for detecting the IC50 dose required to find phenotypic changes in tubulin polymerization. First, the optimal amount of time required to detect tubulin polymerization inhibition of modified HeLa cells treated with colchicine was determined by doing time-lapse imaging for 16 hours. This showed that the tubulin polymerization inhibition process is fast and can be detected in the first three hours of treatment (data not shown). The Haralick homogeneity values of cells treated for three hours were obtained with the three hits compounds and colchicine in a concentration range between 1 nM and 4 uM. The validation showed that the three compounds can inhibit tubulin polymerization (Figure 3A). However, it was also found that the three compounds have different potencies (KX2-391 > ON-01910 > HMN-214) (Figure 3B) and that colchicine inhibits tubulin polymerization at lower doses than the three compounds.

**Figure 3.**
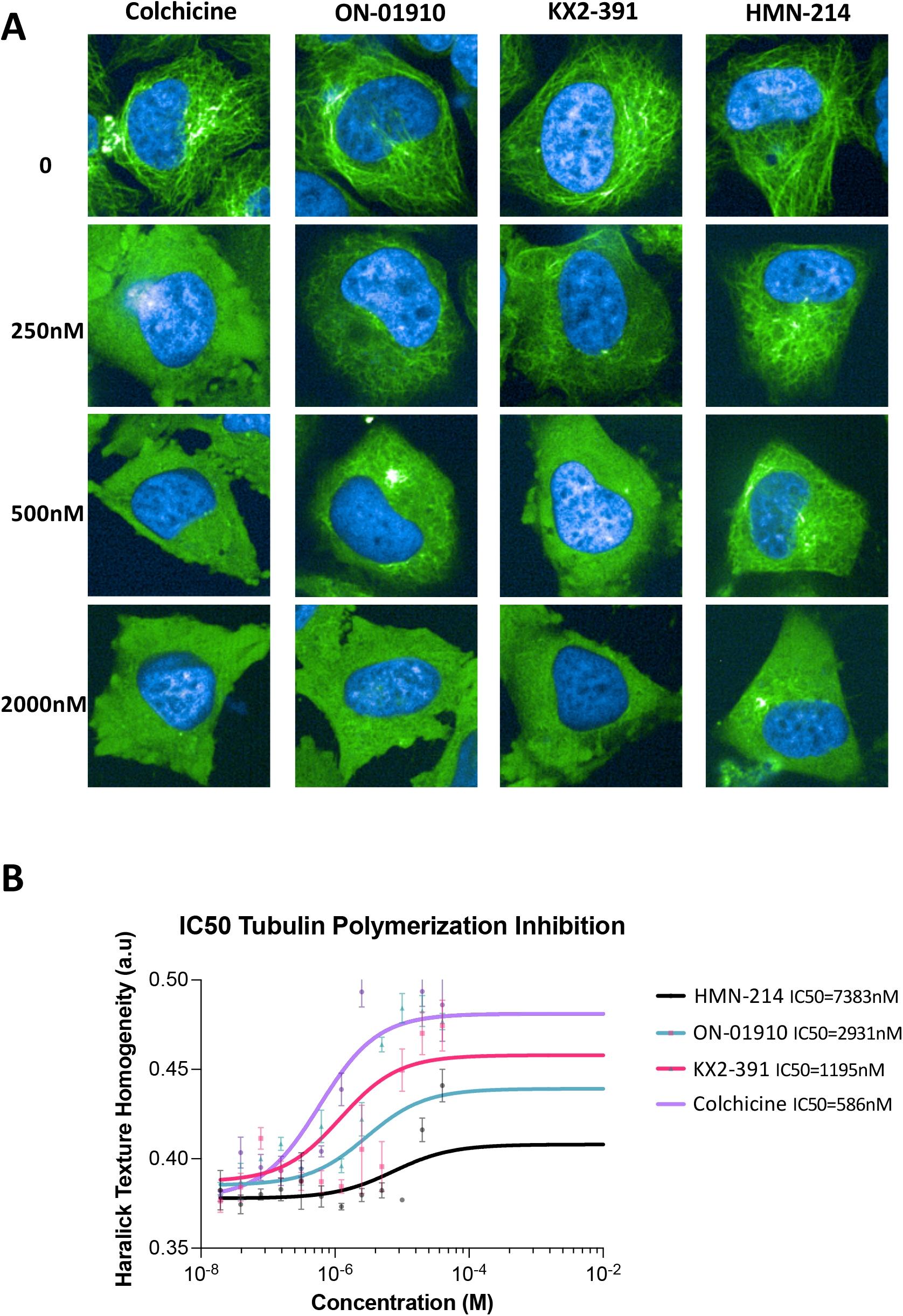
Validation of ON-01910, KX2-391, and HMN-214 as tubulin polymerization inhibitors. (A) Representative images of cells treated with three kinase inhibitors identified as potential tubulin polymerization inhibitors. The cells were treated in triplicates in a range of concentrations between 1 nM and 4000 nM. (B) The Haralick texture homogeneity values of cells treated with the three compounds (1 nM to 4000 nM) was used to determine the phenotypic half maximal tubulin polymerization inhibitory concentration (IC50). Each data point is n = 3; this data is representative of three independent experiments.

Next, it was evaluated how rapidly ON-01910, KX2-391, and colchicine affected tubulin polymerization by tracking treated cells over a period of three hours by acquiring images in the two fluorescent channels every three minutes. After analyzing the time-lapsed images with the routine described in Figure 1A, the dynamic Haralick homogeneity values (Figure 4) were obtained. It was confirmed that the three compounds inhibit tubulin polymerization rapidly, and the changes can be detected in the first 20 minutes of treatment. The results indicate that treatment with colchicine yields the highest change in Haralick texture homogeneity, followed by KX2-391 and then ON-01910. In the control group, the cells maintain the structure of the tubulin and continue to divide; however, for all the treatment groups, the tubulin structures rapidly disappear (Supplementary Video 1).

**Figure 4.**
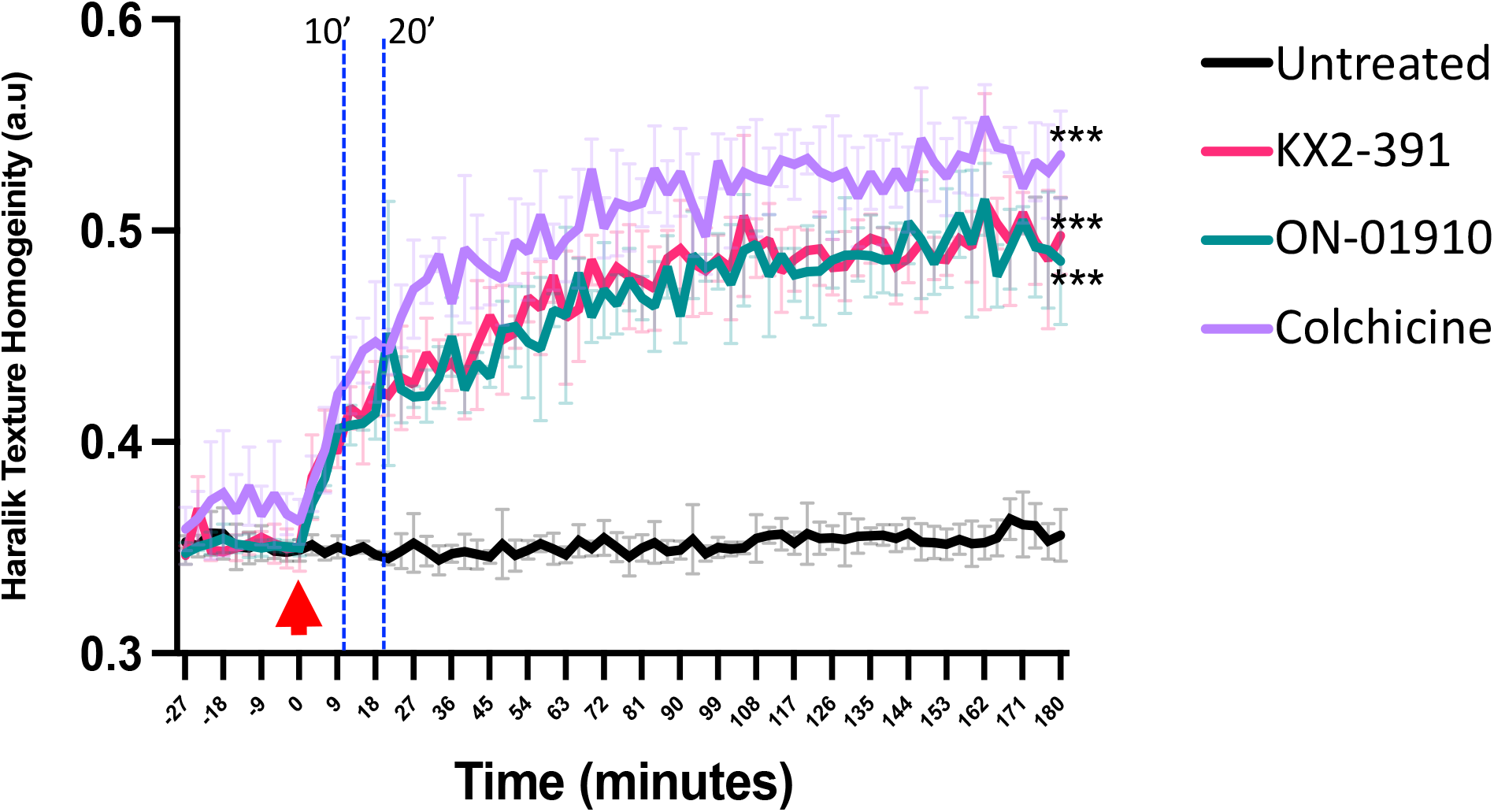
Live cell tracking of the inhibition of tubulin polymerization by detecting changes in the Haralick homogeneity values. HeLa cells with endogenous tagging of β-tubulin and histone 1 were treated with ON-01910, KX2-391, and colchicine as a positive control to detect the tubulin polymerization inhibition over a period of three hours. Imaging capture (every three minutes) started 30 minutes before adding the compounds and continued for a total of three hours. The red arrow indicates the start of the compound treatment. The two vertical dashed lines indicate the 10- and 20-minute timepoints. Differences between drug treatments vs. untreated control were analyzed by two-way ANOVA, and *p* < 0.05 was considered statistically significant. *** *p* < 0.001; error bars represent ± *SD* of triplicate wells. This figure is representative of three independent experiments.

### Molecular docking of kinase inhibitors that inhibit tubulin polymerization

Another aim was to evaluate the potential interaction of the hit compounds (ON-01910, KX2-391, and HMN-214) with β-tubulin and to confirm that the effect of tubulin polymerization inhibition is achieved by directly blocking tubulin function. We evaluated the potential interaction of these compounds with β-tubulin using molecular docking, which is used to predict the predominant binding mode of a ligand with a protein of known three-dimensional structure [18].

To test the approach, molecular docking with known binders of β-tubulin, colchicine, vinblastine, vincristine and paclitaxel was performed using the three-dimensional structure of β-tubulin from *bos taurus*, which has 100% sequence identity with human β-tubulin (TUBB). The predicted interactions match the location of experimental x-ray crystallography structures (Figure 5A). Comparing the docking results of the known binders of β-tubulin, it is evident that colchicine and vinblastine interact with the colchicine binding site, while vincristine and paclitaxel interact with alternate binding sites. Following this validation, a molecular docking prediction was performed of the interaction of the three kinase inhibitors: ON-01910, KX2-391, and HMN-214 with β-tubulin. The analysis suggests that these compounds all interact with the colchicine binding site (Figure 5B) and that multiple amino acids that surround the binding site of colchicine are also present in the predicted binding site of the three kinase inhibitors (Figure 5C). The molecular docking software ranks the binding modes according to the Vina score, an empirical scoring function that assesses the contributions of a variety of different factors and determines the affinity of the protein-ligand interaction [19]. Based on the rankings, the smaller the value (larger negative number) of the Vina score, the more accurate the predicting interaction. Based on the docking results for colchicine, ON-01910, HMN-214, and KX2-391, the Vina scores were -7.0, -7.0, -8.6, and -8.0, respectively.

**Figure 5.**
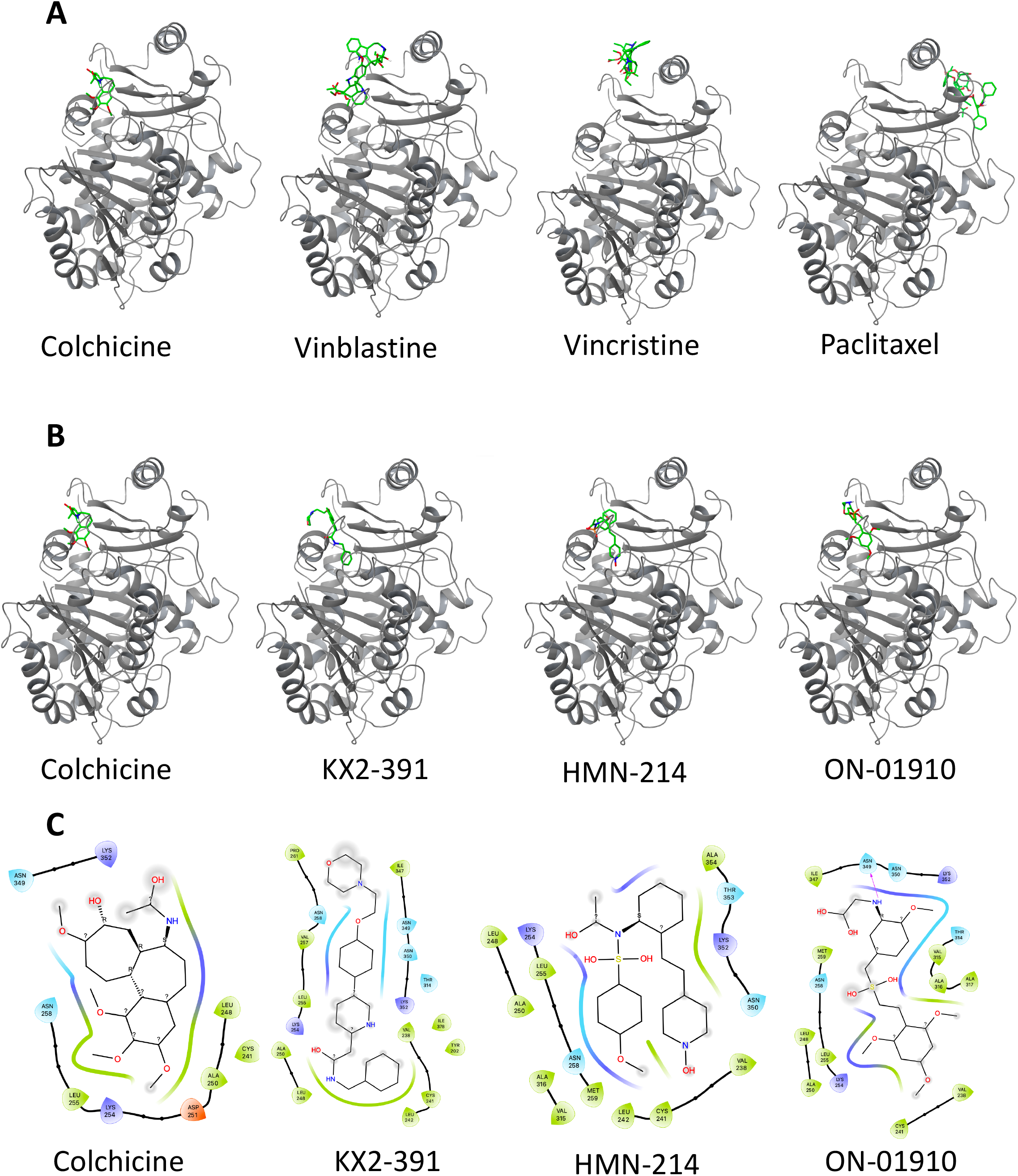
Molecular docking of kinase inhibitors that inhibit tubulin polymerization. (A) Molecular docking identification of the 3D binding site in β-tubulin of known drugs that inhibit tubulin polymerization (colchicine, vinblastine, vincristine) or a drug that can stabilize microtubules (paclitaxel). These predictions were made with CB-Dock 2 web server. (B) Comparison of the β-tubulin predicted binding site of kinase inhibitors (KX2-391, HMN-214, ON-01910) that can inhibit tubulin polymerization against the binding site of Colchicine. (C) Maestro software was used for the Identification of the amino acids surrounding the β-tubulin binding site of colchicine, KX2-391, HMN-214, and ON-01910.

## DISCUSSION

Microtubules play a crucial role in several biological processes, including cellular shape maintenance, cell motility, and mitosis [20]. Due to the clinical efficacy of some microtubule-targeting medicines for treating cancer, microtubules continue to be promising targets for developing novel anti-cancer drugs [21].

This work describes a simplified process to identify tubulin polymerization inhibitors with live cells. The method relies on the use of a CRISPR-modified cell line that contains fluorescent proteins tagged to endogenous β-tubulin and histone 1. This cell line allows the direct use of high-content imaging analysis for cellular segmentation and microtubule network detection without requiring cell fixing or immunostaining. With this method, the three compounds (ON-01910, KX2-391, and HMN-214) were identified in a group of 429 kinase inhibitors as tubulin polymerization inhibitors. Previous reports that used alternate methodologies validate these findings for two of these compounds. ON-01910 was reported to show tubulin destabilization activity after comparing changes in cells with genome-wide upregulation/downregulation via a CRISPR activation/inhibition approach and treated with this compound [22]. In addition, KX2-391 was initially developed as a low-nanomolar inhibitor of Src tyrosine kinase for cancer treatment; however, further studies confirmed that at higher concentrations, KX2-391 has tubulin polymerization inhibition activity [23]. KX2-391 went to clinical trials as a topical tubulin polymerization inhibitor for the treatment of actinic keratosis and received FDA approval in December 2020.

Two of the three compounds identified in this study (ON-01910 and KX2-391) are potent (< 10 nM IC50) kinase inhibitors of their respective target, while HMN-214 is an inactive prodrug that requires liver metabolic transformation before becoming a kinase inhibitor [24]. The capability of these compounds to inhibit tubulin polymerization seems to be independent of their function for inhibiting their kinase target. This assumption is suggested by showing that their capability to inhibit tubulin polymerization only occurs at high concentrations (> 250 nM). In addition, HMN-214 does not have kinase inhibitory activity before going through liver transformation but can inhibit tubulin polymerization at high concentrations.

The molecular docking analysis indicates that these three compounds can interact directly with the colchicine binding site of β-tubulin, thus suggesting a similar mechanism of action for inhibiting tubulin polymerization. The molecular docking was done using CB-Dock version 2, a cavity detection-guided protein-ligand blind docking web server [25]. The current version of CB-Dock 2 has a calculated 85.9% prediction accuracy [25], thus indicating that there is a high probability that the right binding pocket was identified.

This study validates the simple use of a CRISPR-edited cell line for identifying tubulin polymerization inhibitors via high-content imaging analysis. This method is considerably cheaper and less laborious than previous methods due to the elimination of the required steps for cellular fixing and immunostaining. Additionally, this method allows live cell analysis of tubulin polymerization for evaluating kinetic changes. We foresee the use of cell lines with endogenous tagging of tubulin with fluorescence proteins for screening larger and diverse libraries for discovering novel tubulin polymerization inhibitors. We also anticipate that this type of cell lines could be good models to validate novel tubulin polymerization inhibitors discovered by virtual compound screening.

## ACKNOWLEDGEMENTS

This work was supported in part by the National Institute of Health Grants R03DK105267 and R21HG012241.

## AUTHOR CONTRIBUTIONS

HK performed cell culture, drug treatments, and molecular docking and wrote the paper. CAB performed high-content imaging. JG performed automated kinase inhibitor library screening. BO performed cell culture, cell sorting, and single clone cell isolation. OPL conceived the project, designed the experiments, performed high-content imaging, analyzed the data, and wrote the paper.

## SUPPLEMENTARY VIDEO LEGENDS

Supplementary Video 1. Evaluation of the capacity of Colchicine, KX2-391, and ON-01910 for inhibiting tubulin polymerization. Time-lapse confocal microscopy (one image every 3 minutes for 180 minutes) of the HeLa cell line with nuclear (H1-mTagBFP2), and microtubule (β tubulin-mClover3), labeling is shown. The compounds were added 30 min after the start of image acquisition. This time-lapse is representative of three independent experiments.

## REFERENCES

1. Mattiuzzi C, Lippi G: Current Cancer Epidemiology. J Epidemiol Glob Health 2019, 9:217–222.

2. Biemar F, Foti M: Global progress against cancer-challenges and opportunities. Cancer Biol Med 2013, 10:183–186.

3. Debela DT, Muzazu SG, Heraro KD, Ndalama MT, Mesele BW, Haile DC, Kitui SK, Manyazewal T: New approaches and procedures for cancer treatment: Current perspectives. SAGE Open Med 2021, 9:20503121211034366.

4. Haider K, Rahaman S, Yar MS, Kamal A: Tubulin inhibitors as novel anticancer agents: an overview on patents (2013-2018). Expert Opin Ther Pat 2019, 29:623–641.

5. Kaur R, Kaur G, Gill RK, Soni R, Bariwal J: Recent developments in tubulin polymerization inhibitors: An overview. Eur J Med Chem 2014, 87:89–124.

6. Borisy G, Heald R, Howard J, Janke C, Musacchio A, Nogales E: Microtubules: 50 years on from the discovery of tubulin. Nat Rev Mol Cell Biol 2016, 17:322–328.

7. McLoughlin EC, O’Boyle NM: Colchicine-Binding Site Inhibitors from Chemistry to Clinic: A Review. Pharmaceuticals (Basel) 2020, 13.

8. Wattanathamsan O, Pongrakhananon V: Post-translational modifications of tubulin: their role in cancers and the regulation of signaling molecules. Cancer Gene Ther 2021.

9. Harrison MR, Holen KD, Liu G: Beyond taxanes: a review of novel agents that target mitotic tubulin and microtubules, kinases, and kinesins. Clin Adv Hematol Oncol 2009, 7:54–64.

10. Lu Y, Chen J, Xiao M, Li W, Miller DD: An overview of tubulin inhibitors that interact with the colchicine binding site. Pharm Res 2012, 29:2943–2971.

11. Dumontet C, Jordan MA: Microtubule-binding agents: a dynamic field of cancer therapeutics. Nat Rev Drug Discov 2010, 9:790–803.

12. Mukhtar E, Adhami VM, Mukhtar H: Targeting microtubules by natural agents for cancer therapy. Mol Cancer Ther 2014, 13:275–284.

13. Zhu T, Wang SH, Li D, Wang SY, Liu X, Song J, Wang YT, Zhang SY: Progress of tubulin polymerization activity detection methods. Bioorg Med Chem Lett 2021, 37:127698.

14. Wang Z, Chen J, Wang J, Ahn S, Li CM, Lu Y, Loveless VS, Dalton JT, Miller DD, Li W: Novel tubulin polymerization inhibitors overcome multidrug resistance and reduce melanoma lung metastasis. Pharm Res 2012, 29:3040–3052.

15. Perez-Leal O, Nixon-Abell J, Barrero CA, Gordon JC, Oesterling J, Rico MC: Multiplex Gene Tagging with CRISPR-Cas9 for Live-Cell Microscopy and Application to Study the Role of SARS-CoV-2 Proteins in Autophagy, Mitochondrial Dynamics, and Cell Growth. CRISPR J 2021, 4:854–871.

16. Arnst KE, Banerjee S, Chen H, Deng S, Hwang DJ, Li W, Miller DD: Current advances of tubulin inhibitors as dual acting small molecules for cancer therapy. Med Res Rev 2019, 39:1398–1426.

17. Brynolfsson P, Nilsson D, Torheim T, Asklund T, Karlsson CT, Trygg J, Nyholm T, Garpebring A: Haralick texture features from apparent diffusion coefficient (ADC) MRI images depend on imaging and pre-processing parameters. Sci Rep 2017, 7:4041.

18. Meng XY, Zhang HX, Mezei M, Cui M: Molecular docking: a powerful approach for structure-based drug discovery. Curr Comput Aided Drug Des 2011, 7:146–157.

19. Quiroga R, Villarreal MA: Vinardo: A Scoring Function Based on Autodock Vina Improves Scoring, Docking, and Virtual Screening. PLoS One 2016, 11:e0155183.

20. Ilan Y: Microtubules: From understanding their dynamics to using them as potential therapeutic targets. J Cell Physiol 2019, 234:7923–7937.

21. Kuppens IE: Current state of the art of new tubulin inhibitors in the clinic. Curr Clin Pharmacol 2006, 1:57–70.

22. Jost M, Chen Y, Gilbert LA, Horlbeck MA, Krenning L, Menchon G, Rai A, Cho MY, Stern JJ, Prota AE, et al: Combined CRISPRi/a-Based Chemical Genetic Screens Reveal that Rigosertib Is a Microtubule-Destabilizing Agent. Mol Cell 2017, 68:210–223 e216.

23. Smolinski MP, Bu Y, Clements J, Gelman IH, Hegab T, Cutler DL, Fang JWS, Fetterly G, Kwan R, Barnett A, et al: Discovery of Novel Dual Mechanism of Action Src Signaling and Tubulin Polymerization Inhibitors (KX2-391 and KX2-361). J Med Chem 2018, 61:4704–4719.

24. Takagi M, Honmura T, Watanabe S, Yamaguchi R, Nogawa M, Nishimura I, Katoh F, Matsuda M, Hidaka H: In vivo antitumor activity of a novel sulfonamide, HMN-214, against human tumor xenografts in mice and the spectrum of cytotoxicity of its active metabolite, HMN-176. Invest New Drugs 2003, 21:387–399.

25. Liu Y, Yang X, Gan J, Chen S, Xiao ZX, Cao Y: CB-Dock2: improved protein-ligand blind docking by integrating cavity detection, docking and homologous template fitting. Nucleic Acids Res 2022.

